# Selection on non-antigenic gene segments of seasonal influenza A virus and its impact on adaptive evolution

**DOI:** 10.1101/166082

**Authors:** Jayna Raghwani, Robin Thompson, Katia Koelle

## Abstract

Most studies on seasonal influenza A/H3N2 virus adaptation have focused on the main antigenic gene, haemagglutinin. However, there is increasing evidence that the genome-wide genetic background of novel antigenic variants can influence these variants’ emergence probabilities and impact their patterns of dominance in the population. This suggests that non-antigenic genes may be important in shaping the viral evolutionary dynamics. To better understand the role of selection on non-antigenic genes in the adaptive evolution of seasonal influenza viruses, we here develop a simple population genetic model that considers a virus with one antigenic and one non-antigenic gene segment. By simulating this model under different regimes of selection and reassortment, we find that the empirical patterns of lineage turnover for the antigenic and non-antigenic gene segments are best captured when there is both limited viral coinfection and selection operating on both gene segments. In contrast, under a scenario of only neutral evolution in the non-antigenic gene segment, we see persistence of multiple lineages for long periods of time in that segment, which is not compatible with the observed molecular evolutionary patterns. Further, we find that reassortment, occurring in coinfected individuals, can increase the speed of viral adaptive evolution by primarily reducing selective interference and genetic linkage effects mediated by the non-antigenic gene segment. Together, these findings suggest that, for influenza, with 6 internal or non-antigenic gene segments, the evolutionary dynamics of novel antigenic variants are likely to be influenced by the genome-wide genetic background as a result of linked selection among both beneficial and deleterious mutations.

## INTRODUCTION

Seasonal influenza is a major infectious disease that causes 3 to 5 million worldwide cases of severe illness and 250,000 to 500,000 deaths each year in humans (1). Of the currently circulating flu viruses, influenza A subtype H3N2 is the predominant virus contributing to these morbidity and mortality estimates. This virus is known to rapidly evolve, particularly antigenically (2), enabling it to perpetually evade herd immunity and re-infect individuals in the population. Consequently, there has been great interest in understanding how this virus evolves antigenically, especially with respect to its main antigenic gene, haemagglutinin (HA). In particular, these investigations have focused on identifying key sites involved in viral antigenicity (3-6), which has provided compelling evidence of immune-mediated selection acting upon HA.

However, the limited standing genetic diversity observed for HA has been difficult to reconcile based on recurrent positive selection alone, since the high virus mutation rate and the presence of strong diversifying selection predicts a large antigenic repertoire over time (7). The observed low-level genetic diversity of the HA is reflected in its spindly, ladder-like phylogeny, which indicates that only a single viral lineage persists over time. Genetic variants belonging to this persisting lineage have been characterized antigenically, indicating that every two to eight years a major antigenic change occurs that necessitates the updating of components of the seasonal influenza vaccine (6, 8, 9). Phylodynamic models have proven to be invaluable to understanding how host immunity and viral evolution can lead to these interesting phenomena of a spindly phylogeny and a single major circulating antigenic variant dominating global infection dynamics (7, 10-12). While these models differ in their specific explanations of what processes shape this restricted antigenic evolution of influenza A/H3N2, they in general have had to either impose strong among-strain competition for susceptible hosts (7, 11, 12) and/or limit the antigenic mutation rate (10, 12). More recent work on the molecular evolution of the HA indicates that clonal interference and background selection are also important determinants of the adaptive dynamics of the HA (13-17).

While it is clear that the evolution of HA is a key component of influenza A/H3N2’s adaptive evolution, the role of other gene segments, in particular those that encode internal proteins, is less well understood. There is a small but growing number of studies that indicate that selection also acts on viral phenotypes beyond antibody-mediated immune escape. For example, the appearance and dominance of the CA04 antigenic lineage is attributed in part to the increased replicative fitness and virulence conferred by two amino-acid substitutions in the polymerase acidic (PA) gene segment (18). There is also evidence that cytotoxic T-lymphocyte (CTL) immune pressure can exert selection pressure on influenza A virus. Specifically, recent work has shown that adaptive substitutions in the nucleoprotein (NP) gene predominantly occur at T-cell epitopes (19, 20).

Interestingly, the genetic diversity of internal or non-antigenic genes in influenza A/H3N2 virus is also limited, although to a lesser extent than for HA (21). One explanation for this observation is that these gene segments are in strong linkage with HA, which means that any evolutionary force that reduces genetic diversity of the HA (e.g., selective sweeps and genetic bottlenecks) will also similarly impact the rest of the virus genome. However, whole-genome analyses of seasonal influenza A viruses indicate that reassortment is relatively frequent, with each gene segment having somewhat of a distinctive evolutionary history (21-25). Estimated differences in the times to most recent common ancestor (TMRCAs) across the genome can also exceed six years (21), which is inconsistent with strong linkage effects solely shaping the genetic diversity patterns of this virus. An alternative explanation for the limited genetic diversity of non-antigenic gene segments is selection. Although there are several distinct models that can generate the restricted diversity of HA by invoking selection (10-12, 14, 15, 26), there has been very little consideration of whether selection also contributes to shaping the evolutionary dynamics of non-antigenic genes.

Here we evaluate the importance of selection on non-antigenic gene segments in the adaptive evolution of seasonal influenza A/H3N2 by analyzing the evolutionary dynamics of the viral genome and using a population genetic model to determine the critical processes that can reproduce features of these observed evolutionary dynamics. The main questions we address are whether selection on non-antigenic gene segments impact the evolutionary dynamics of the non-antigenic gene segments themselves, and through linkage effects, the antigenic gene segments. Instead of examining the complexity of 8 distinct gene segments, we simplify our model by considering a virus that contains only two gene segments, corresponding to one antigenic gene (e.g., HA) and one non-antigenic gene (e.g., PA). By simulating the model such that lineages can be traced back in time, we examine the patterns of genetic diversity of the virus across different assumptions of selection and reassortment. We find that selective effects on both gene segments and limited reassortment (via limited coinfection rates) are necessary to capture the key TMRCA patterns of influenza A/H3N2 virus genome. Furthermore, we find that the rate of adaptive evolution of the virus increases under this evolutionary regime, which is predominantly a result of reassortment reducing interference effects contributed by the non-antigenic gene segment.

## MATERIALS AND METHODS

### A) Evolutionary dynamics of seasonal influenza A/H3N2 virus genome

To characterize the evolutionary dynamics of A/H3N2, we used a published global whole-genome dataset of viruses sampled from 1977 to 2009 (*n* = 676) (27). Time-scaled trees were estimated with BEAST v1.8 (28) by employing a relaxed uncorrelated log-normal distributed molecular clock (29), a codon-structured nucleotide substitutional model (30), and a Bayesian Skygrid coalescent prior (31). Two independent chains of 200 million steps were executed for each of the eight gene segments to ensure that adequate mixing and stationarity had been achieved. The posterior tree distribution for each segment was further examined with PACT (32), which infers the times to the most recent common ancestor (TMRCAs) across the entire evolutionary history at regular intervals. To quantify and visualize patterns of genetic diversity in each segment, mean TMRCAs over time were plotted using the R package ggplot2 (33) and genealogical trees were plotted with ggtree (34).

### B) Phylodynamic model of infection and coinfection

To explore the evolutionary processes underlying the empirical patterns of TMRCA observed for the influenza A/H3N2 virus genome, we formulated a simple population genetic model with a constant number of *N* = 1000 infected individuals. In the model, individuals were either infected with a single virus (*I*_s_) or coinfected with two viruses (*I*_co_). We did not consider coinfection with more than two viruses. The virus genome consisted of one antigenic segment and one non-antigenic segment.

We simulated the infected population of hosts over time using a modified Moran model. Specifically, we allowed for two types of infection events: ‘infection/recovery’ events and coinfection events. When an ‘infection/recovery’ event occurred, an infected individual (*I*_s_ or *I*_co_) was chosen to generate a new singly-infected individual. If it was a coinfected individual generating the new infection, that individual transmitted either of the viruses he was infected with or a reassortant virus (Figure 1A). We assumed an equal probability of each viral gene segment being transmitted, such that a reassortant strain was transmitted 50% of the time. At the same time as the generation of the new infection occurred, recovery of a randomly chosen infected individual (*I*_s_ or *I*_co_) also occurred. If a coinfected individual was chosen to recover, he cleared both infecting viral strains. Because infection events were always offset by recovery events, as is traditional in Moran models where a ‘birth’ is always offset by a ‘death’, the total number of infected individuals in the population remained constant (Figure 1B). Infection/recovery events occurred at a rate of *α* = 0.25 per capita per day, reflecting a typical duration of influenza infection of approximately 4 days (35).

**Figure 1:**
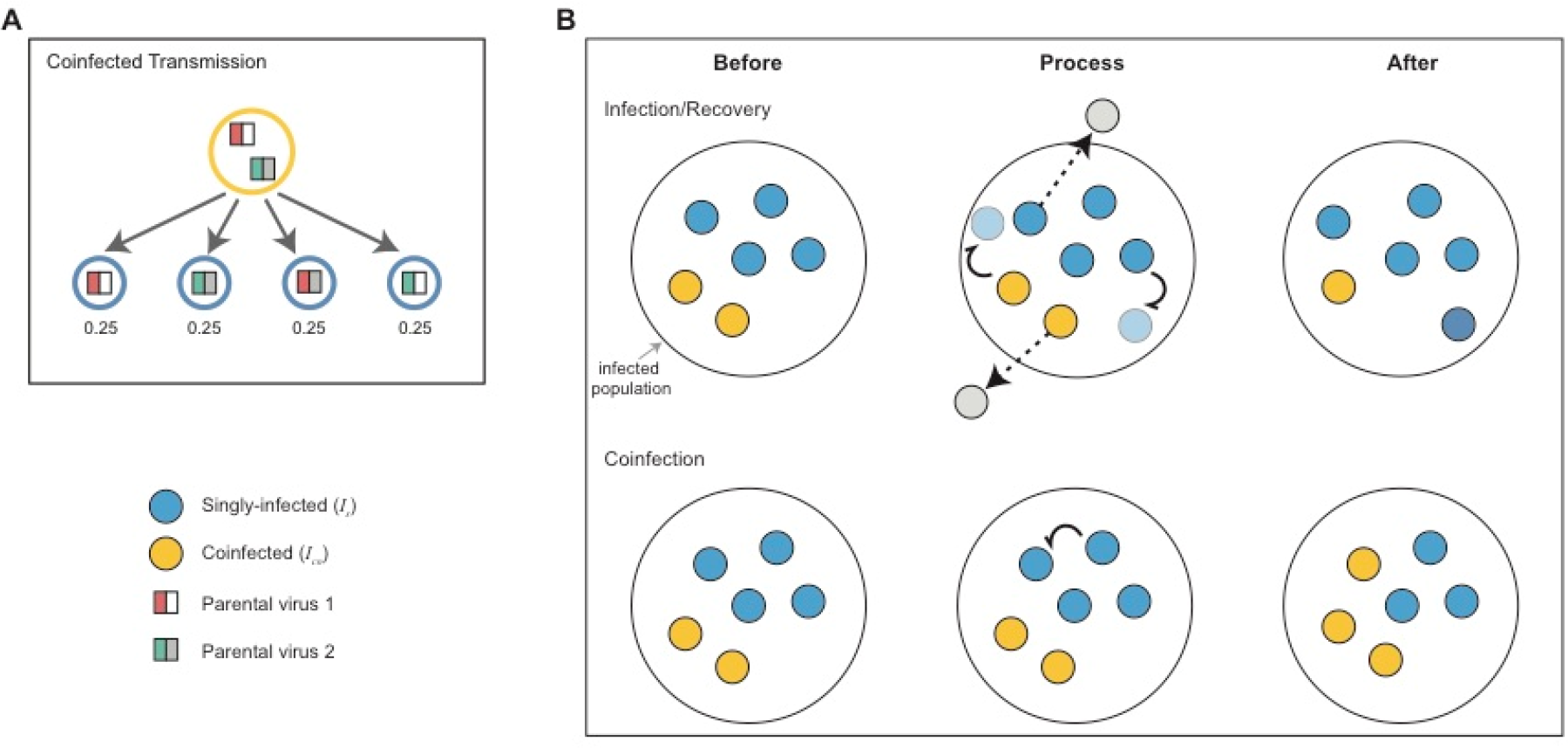
Schematic of the events in the population genetic model. (A) Infection transmission by a coinfected individual. When a coinfected individual transmits, each gene segment is randomly chosen from the two viral strains present in that individual. Consequently, there is an equal probability of transmitting a non-reassortant strain (two strains on the left) as there is of transmitting a reassortant strain (two strains on the right). (B) Schematic of the main events: infection/recovery and coinfection. Infection and recovery are coupled, such that the infected population remains constant. Upon infection (indicated by curly arrows), singly-infected individuals (blue circles) and coinfected individuals (yellow circles) generate new singly-infected individuals. Recovery of singly-infected and coinfected individuals removes them from the population (denoted by dashed arrows). Here, two infection/recovery events are shown that occur in the same *τ* time step. Coinfection events occur when a singly-infected individual infects another singly-infected individual. This results in a new coinfected individual in the infected population, carrying two viral strains. Coinfection events result in an increase in the number of coinfected individuals in the population and a decrease in the number of singly-infected individuals.

Coinfection events were marked by singly-infected individuals infecting other singly-infected individuals (Figure 1B). Coinfection events occurred from singly infected individuals at a per capita rate of *β* = 0.0125 per day. This corresponds to a coinfection level of approximately 5% of the total infected population at equilibrium (see Text S1). Ascertaining an empirical coinfection rate for influenza A/H3N2 viruses in general, or at the within-subtype level, is very difficult, since the low circulating viral diversity is likely to limit our ability to distinguish between independent infecting viral strains. Nevertheless, the number of influenza coinfections can be estimated when viral strains involved belong to either different subtypes or types (e.g. A/H3N2 and A/H1N1 or influenza A and B viruses, respectively) (36, 37). These types of coinfection have been known to occur between 1-2% in sampled influenza A viral infections (36-38). We set the level of coinfection in our model slightly higher than these empirical estimates, at ~5%, to reflect that these empirical estimates between different subtypes or types are likely underestimates.

### Evolution of the antigenic and non-antigenic gene segments

We let mutations occur at transmission events, which consist of both ‘infection’ events and ‘coinfection’ events. We let the number of new mutations present in the transmitting virus be Poisson-distributed with mean *U* = 0.1, with each mutation being equally likely to land on the antigenic or the non-antigenic gene segment. We allow the distribution of mutational fitness effects to differ between the two gene segments. Specifically, we assume that 30% of mutations are beneficial and 70% of mutations are deleterious on the antigenic gene segment. On the non-antigenic gene segment, we assume that 5% of mutations are beneficial, 30% of mutations are deleterious, and the remaining 65% of mutations are neutral. A higher proportion of beneficial mutations are assumed in the antigenic gene segment to capture the selective advantage that antigenic mutations are likely to have through evasion of herd immunity. The non-antigenic gene segment is assumed to have a greater proportion of neutral mutations to reflect the observation that internal genes undergo greater neutral evolution than external genes (25). We assume that the fitness effects for beneficial mutations are exponentially distributed with mean 0.03 and that the fitness effects for deleterious mutations are exponentially distributed with mean 0.09. We do not consider lethal mutations. Importantly, the distributions of mutational fitness effects on the antigenic and non-antigenic gene segment capture the salient features of recently determined mutational fitness effects for seasonal influenza A virus (39) (see Figure S1).

Viral fitness is calculated by multiplying fitness values at each site across the genome. Multinomial sampling based on viral fitness is applied at each transmission event to determine which individual will infect (or coinfect) next. For coinfected individuals, we initially determine which virus is transmitted from the two infecting parental viral strains (see Figure 1A) and compute the viral fitness accordingly.

### Tracking lineages over time

The model is implemented in Java using a Gillespie tau-leap algorithm (40) for computational efficiency with a time step *τ* of 0.25 days. Starting from an equilibrium number of singly- and coinfected individuals (Text S1), we run each simulation for 60 years, analysing results only from the last 20 years.

To be able to infer the genealogical history of the viral population, we track in our model who-infected-whom at the level of infected individuals and for each gene segment. A random sample of 100 singly-infected individuals is used to infer the TMRCA of each gene segment at yearly intervals. Viral gene genealogies are reconstructed from the last twenty years of simulation using a random sample of 300 singly-infected individuals. The tracked infection histories are used to determine the first ‘coalescent’ event, which corresponds to finding the two sampled individuals that shared the most recent common ancestor for a given gene segment. Specifically, this process involves tracing back the transmission events from the sampled infections, and establishing the parental virus in common with the most recent transmission time. This procedure is repeated until all sampled and ancestral lineages reach the parental viral infection that represents the most recent common ancestor of the entire sample.

## RESULTS

### Genealogical diversity of seasonal influenza A/H3N2 virus

Figure 2 shows how the genealogical diversity of seasonal influenza A/H3N2 varies over time for each gene segment. We observe that the mean TMRCA of the HA gene segment (1.90 years) is 0.5-1.1 years younger than the other gene segments, indicating that HA experiences the fastest lineage turnover in the virus genome. The maximum TMRCA for this gene segment also does not exceed 5 years. These TMRCA patterns reflect that the HA gene genealogy has a single viral lineage dominating over time (Figure S2). NA is found to have the second lowest mean TMRCA (2.4 years), indicative of slightly longer lineage persistence than HA (Figure S2). The non-antigenic gene segments of A/H3N2 are marked by larger mean TMRCAs and by more extensive variation in genealogical diversity over time, indicating that multiple lineages can co-exist for significant periods, e.g. up to ~7 years in M1 (Figure S2). Together, these observations are compatible with positive selection predominantly acting upon the antigenic genes, most notably the HA.

**Figure 2:**
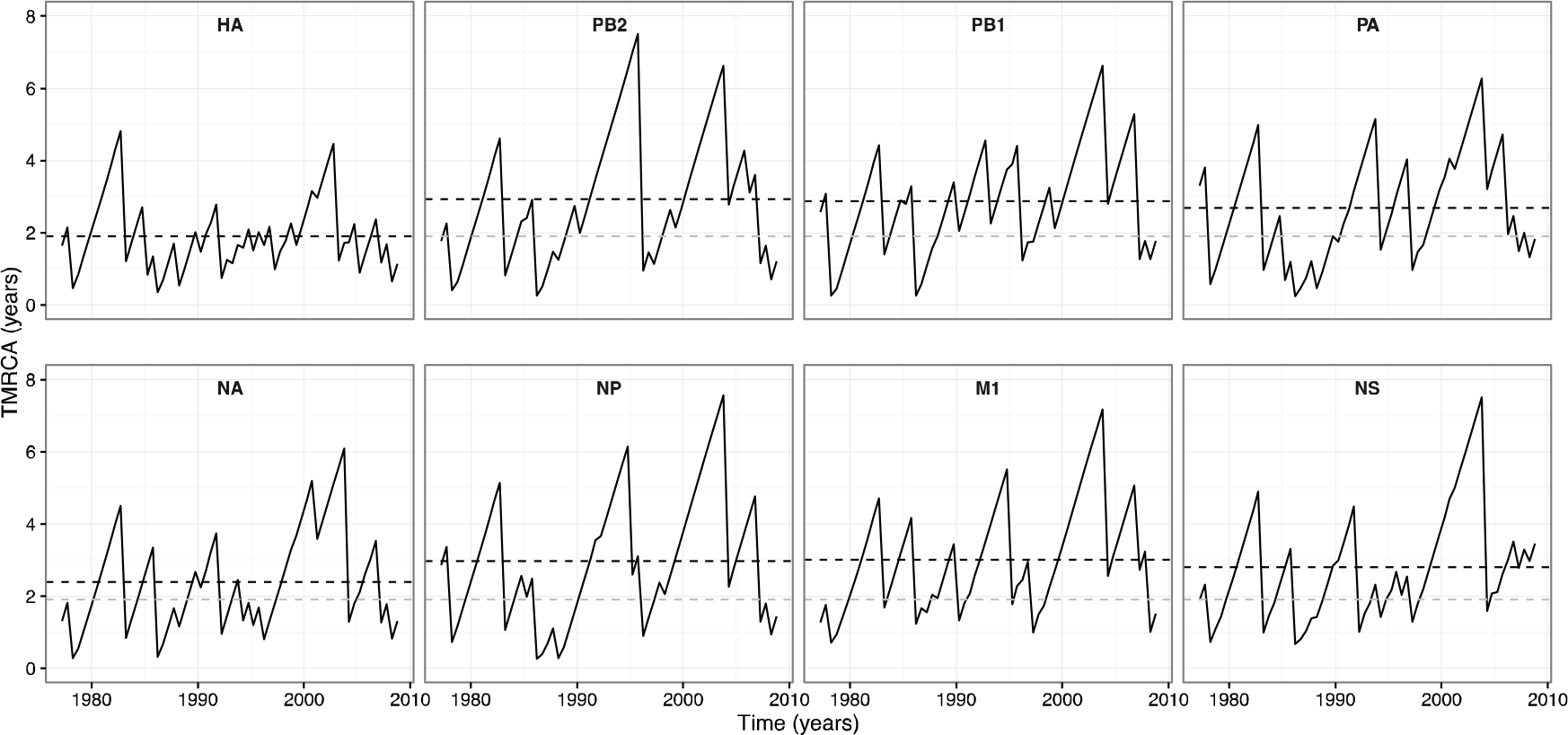
TMRCA through time plots for individual seasonal influenza A/H3N2 gene segments. The mean TMRCA over time is estimated from a posterior tree distribution for each gene segment at 6-month intervals. The black dashed lines indicate the overall mean TMRCA for the focal gene segment in each subplot. The gray dashed lines in the non-HA gene segment subplots show the overall mean TMRCA for the HA.

### Evolutionary dynamics when only antigenic gene segment is under selection

To better understand the patterns of genealogical diversity of the influenza A/H3N2 virus genome, we first simulated the described model under the assumption that the adaptive evolution of the virus is restricted to the antigenic gene segment, with all mutations on the non-antigenic gene segment assumed to be neutral. Further, coinfection was not permitted (*β* = 0). Figure 3 shows results from a representative simulation. Viruses with higher fitness constantly emerge and become dominant in the population over time (Figure 3A). At any given time, significant fitness variation is present in the population, with lower fitness viruses able to persist in the population over extended periods (Figure 3A).

**Figure 3:**
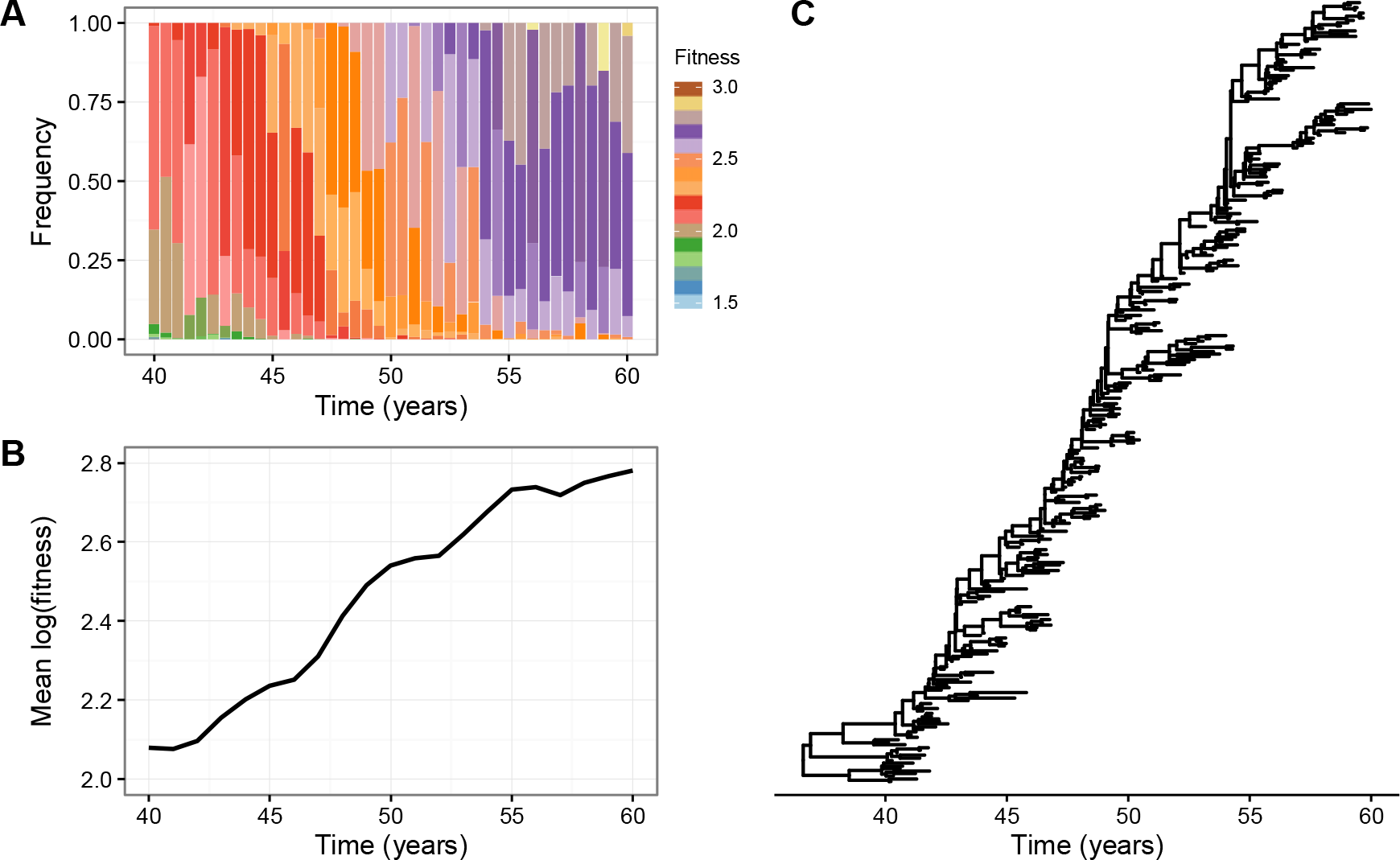
Adaptive evolution in the absence of coinfection and when selection acts only on the antigenic gene segment. (A) The distribution of fitness in the viral population over time. (B) Mean (log) fitness of the virus population over time. (C) Gene genealogy reconstructed from model simulation by sampling 300 singly-infected individuals over 20 years, following a burn-in of 40 years.

Interestingly, the simulated viral population evolves in a punctuated manner, as indicated by the change in mean (log) population fitness over time (Figure 3B). This suggests that the tempo of adaptive evolution varies over time. If we consider only beneficial mutations on the antigenic gene segment, we instead observe a smooth and continual increase in the mean population fitness over time (Figure S3). These results indicate that more complex evolutionary dynamics can emerge with a broader distribution of fitness effects. Lastly, consistent with previous studies (15-17), this evolutionary regime where clonal interference and background selection are present reproduces HA’s spindly phylogeny (Figure 3C).

Under the model with both positive and negative fitness effects on only the antigenic gene segment, no significant changes in the mean TMRCA of the antigenic gene segment occur with increasing levels of coinfection (Figure 4A). In contrast, the mean TMRCA of the non-antigenic gene segment becomes notably greater as the rate of coinfection increases. These results are consistent with reassortment reducing the hitchhiking of the neutrally evolving non-antigenic gene segment with the non-neutrally evolving antigenic gene segment. As a consequence, the non-antigenic gene segment is able to explore more genetic backgrounds, which leads to an increase in its genetic diversity. Expectedly, the rate of adaptive evolution is unaffected by changes in the rate of coinfection (Figure 4B).

**Figure 4:**
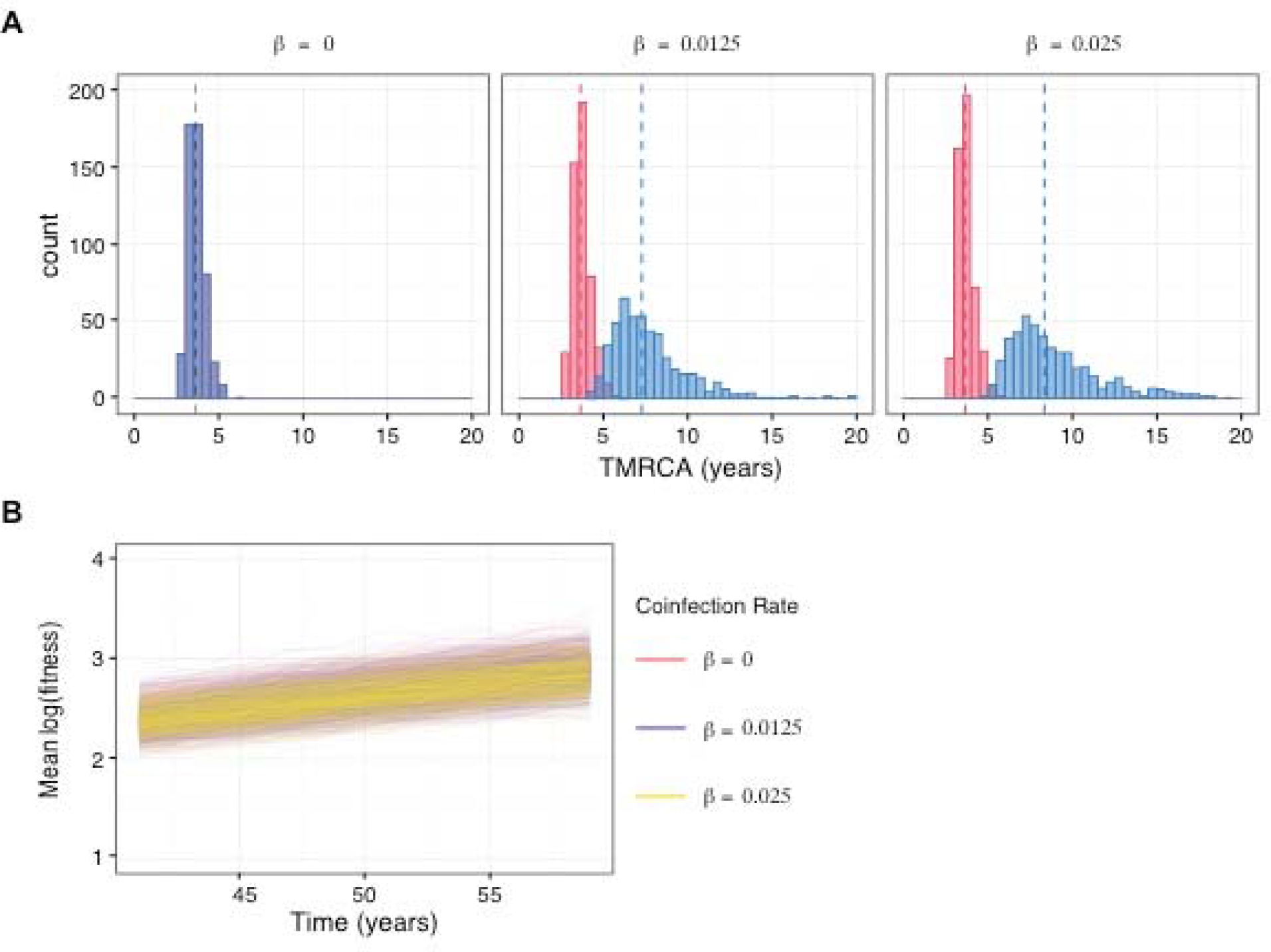
Genealogical diversity and the rate of adaptive evolution at different levels of coinfection when selection acts only on the antigenic gene segment. (A) Distribution of TMRCAs of the antigenic (red) and non-antigenic (blue) gene segments at different levels of coinfection. Three different coinfection levels were considered: 0% *β* = 0), 5% (*β* = 0.0125), and 9% *β* = 0.025). 500 simulations were used to obtain the TMRCA distribution at each of the three coinfection levels. The dashed lines show the mean TMRCA for the focal gene segment in each subplot. (B) Mean (log) fitness of the virus population over time under different coinfection rates.

### Reassortment increases the rate of adaptive evolution when a non-antigenic gene segment is under selection

Next, we examined the behaviour of the model when selection occurs on both gene segments. First, we looked at the changes in mean population fitness and fitness variation over time under increasing coinfection rates (*β* = 0, 0.0125, and 0.025 per day), for the whole virus (Figure 5A), the antigenic gene segment (Figure 5B), and the non-antigenic gene segment (Figure 5C). Strikingly, the rate of virus adaptation is significantly greater in the presence of coinfection than when it is absent (Figure 5A). This phenomenon appears to be primarily driven by the non-antigenic gene segment, which also experiences a notably higher rate of adaptive evolution when coinfection occurs in the population (Figure 5C). In contrast, although coinfection increases the fitness variation of both gene segments (Figure 5B and C), the difference in the rate of adaptive evolution of the antigenic gene segment in the absence versus in the presence of coinfection appears to be slight (Figure 5B).

**Figure 5:**
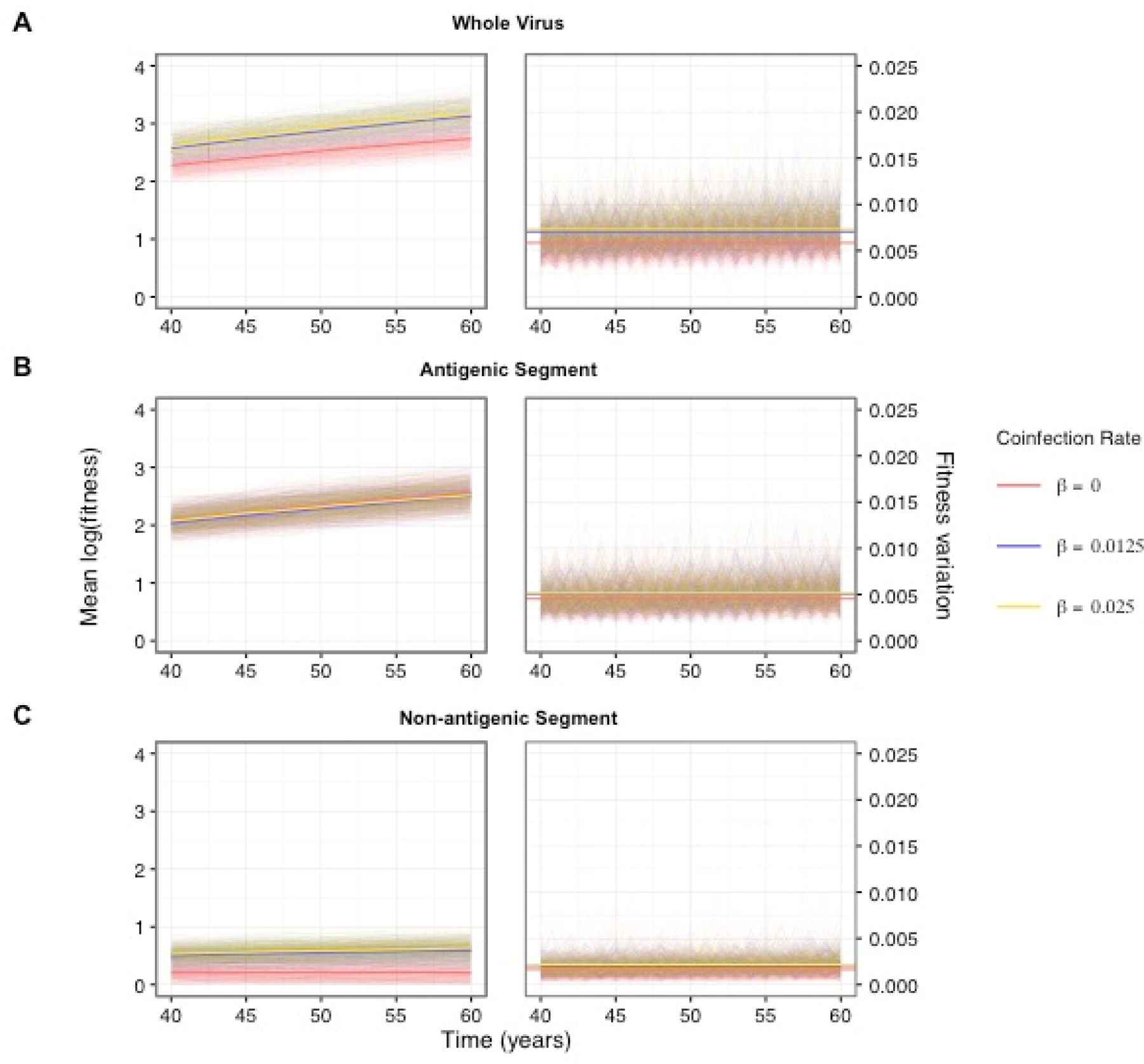
Adaptive evolution of the virus when both gene segments undergo selection at varying levels of coinfection. The mean (log) population fitness and population fitness variation are shown for (A) the whole virus, (B) the antigenic gene segment, and (C) the non-antigenic gene segment for three different coinfection rates, corresponding *β* = 0, *β* = 0.0125, and *β* = 0.025.

Together, these results indicate that when coinfection is absent the non-antigenic gene segment experiences greater selective interference (both among beneficial and deleterious mutations) and genetic hitchhiking than the antigenic gene segment, for the simple reason that there are significantly more mutations with selective effects on the latter. In other words, when there is strong linkage between the two segments, selection on the antigenic gene segment will have a larger impact on the non-antigenic gene segment than the non-antigenic gene segment will have on the antigenic gene segment. As a consequence, while reassortment is expected to reduce linkage effects between both gene segments, larger gains in fitness are more likely for the non-antigenic gene segment as it can explore comparatively more advantageous genetic backgrounds.

These evolutionary dynamics also have an impact on the mean TMRCAs of both gene segments (Figure 6). Although this pattern is more discernible for the non-antigenic gene segment, it does indicate that the antigenic gene segment is influenced to some degree by linkage effects from the non-antigenic gene segment. The larger increase in the mean TMRCA for the non-antigenic gene segment is also consistent with the non-antigenic gene segment experiencing comparatively greater linkage effects than the antigenic gene segment.

**Figure 6:**
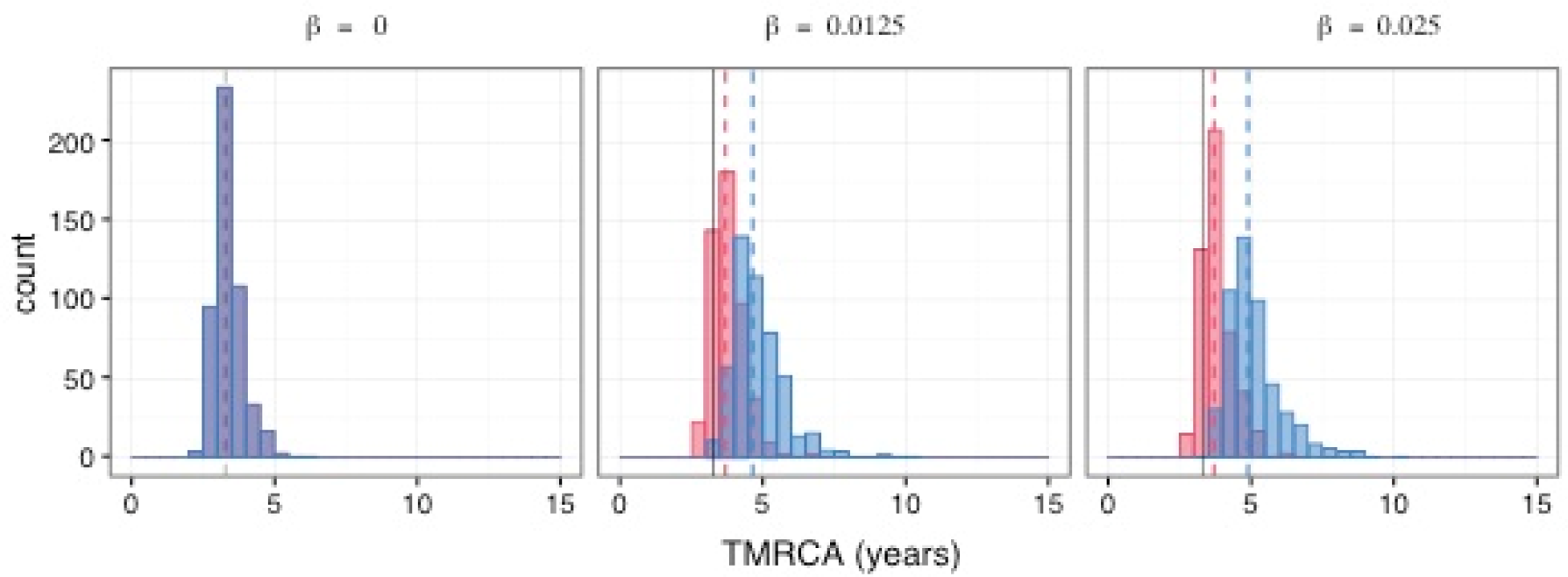
Genealogical diversity of the virus when selection acts on both gene segment at varying levels of coinfection. The distribution of TMRCAs for antigenic (red) and non-antigenic (blue) gene segments for three different coinfection rates: *β* = 0, *β* = 0.0125, and *β* = 0.025. As in Figure 4, 500 simulations were used to obtain each TMRCA distribution at each of the three coinfection levels. The dashed lines show the mean TMRCAs for the each of the two gene segments in each subplot. The solid black lines in the subplots indicates the mean TMRCA of the gene segment when there is no coinfection.

### Empirical genealogical patterns are compatible with selection on both antigenic and non-antigenic gene segments

Figure 7 shows representative gene genealogies for the antigenic and non-antigenic gene segments in the presence of coinfection (*β* = 0.0125) and, in both cases, when selection acts on the antigenic gene segments. The genealogies of the antigenic gene segment when the non-antigenic gene segment evolves either selectively (Figure 7A) or neutrally (Figure 7D) are topologically similar, with a single lineage persisting over time in both cases. In contrast, quite different gene genealogies are observed for the non-antigenic gene segment (Figures 7B and E). Specifically, the non-antigenic gene segment is associated with significantly lower genealogical diversity when it is under selection (Figure 7B) compared to when it is not (Figure 7E). In both cases, however, there is still greater persistence of multiple lineages relative to the antigenic gene segment, indicative of slower population turnover.

**Figure 7:**
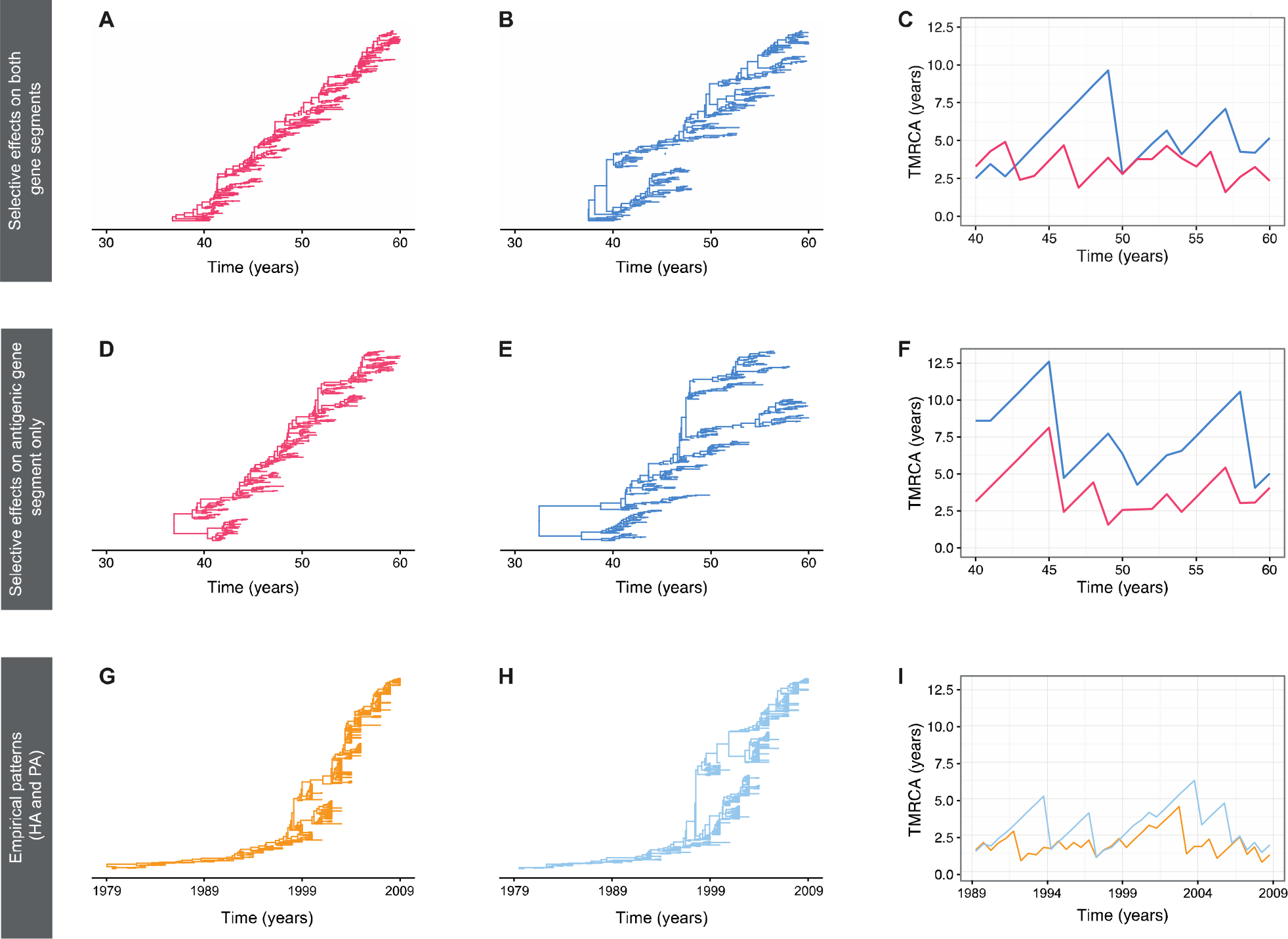
Representative gene genealogies and TMRCA dynamics from simulations when the non-antigenic gene segment evolves either neutrally or under selection. (A-C) Results obtained from model simulations with coinfection and selective effects occurring on both gene segments. (D-F) Results obtained from model simulations with coinfection but no selection on the non-antigenic gene segment. Panels A and D depict the gene genealogies of the antigenic gene segment. Panels B and E depict the gene genealogies of the non-antigenic gene segment. Panels C and F show the TMRCA dynamics of both gene segments over time. The red and blue lines correspond to the antigenic and non-antigenic gene segments, respectively. (G-I) Inferred influenza A/H3N2 MCC phylogenies for the HA gene segment and the PA gene segment, along with their TMRCA dynamics. HA is the dominant antigenic gene segment. PA is a non-antigenic gene segment. The TMRCA dynamics for HA and PA are shown in panel I in orange and light blue lines, respectively.

When comparing TMRCA patterns between the antigenic gene segment and the non-antigenic gene segment, it is notable that when the non-antigenic gene segment evolves neutrally, the common ancestor of the non-antigenic gene segment is consistently older than the antigenic gene segment (Figure 7F). However, when selection affects both gene segments, we note a closer correspondence with the empirical TMRCA dynamics (Figure 7C, compared to Figure 7I)). Specifically, in addition to the antigenic gene segment undergoing more frequent fluctuations in the TMRCA over time compared to the non-antigenic gene segment, the TMRCA of both gene segments can occasionally coincide, which likely indicates a shared common ancestor, perhaps as a result of a genome-wide selective sweep. We further examined this observation by comparing the differences in TMRCA between the gene segments (Figure S4). The higher density around zero years of difference in the TMRCA suggests that the likelihood of sharing a common ancestor is greater when selective effects occur on both gene segments (Figure S4).

### Sensitivity of results to model parameters

#### a) Infected population size

While it is well established that human influenza A/H3N2 virus has a strong seasonal transmission pattern in some populations, we decided to model a constant infected population. This decision was motivated largely by undertaking a simple and standard approach to examine the patterns of viral diversity due to selection, mutation, and reassortment alone. However, given that regions with low-level, constant disease transmission (e.g. the tropics) frequently seed seasonal outbreaks in temperate locales (21, 41-43), the effective population size of global influenza A/H3N2 viruses is expected to be relatively small and constant over time (21, 42). Consequently, the assumption of a constant infected population size is not unreasonable since regions with year-round influenza infection are expected to ultimately shape the overall evolutionary dynamics of the virus. We tested the effects of population size on the evolutionary behavior of the model (Figure S5). Specifically, we ran 100 simulations at each of three population sizes (*N* =1000, *N* = 5000, and *N* =10000), under the model parameterization with both gene segments experiencing positive and negative selection. Notably, similar TMRCA patterns were observed regardless of population size, such that the antigenic gene segment typically had a younger TMRCA compared to the non-antigenic gene segment (Figure S5). However, as we increase the population size, the TMRCA of both gene segments increases, indicating greater lineage persistence in the population. This corroborates a standard expectation from coalescent theory: smaller populations have comparatively more recent common ancestors than larger populations due to stronger effects of genetic drift. Thus, when the infected population size is fixed at N=1000, deterministic and stochastic forces will shape the population’s genetic diversity, both of which have been implicated in the evolutionary dynamics of seasonal influenza A viruses (21). At larger population sizes, we can, however, recover lower TMRCAs when we increase the mean effect size of mutations (results not shown).

#### b) Mutation and coinfection rates

As it is difficult to ascertain the per-genome, per transmission, mutation rate for a two-segment virus, we varied the per-genome per-transmission mutation rate *U* between 0.05 and 0.2 (Figure S6). Overall, these simulations yielded qualitatively similar results: the antigenic gene segment had a younger TMRCA than the non-antigenic gene segment. Interestingly, at higher mutation rates, mean TMRCAs for the non-antigenic gene segment were appreciably smaller and mean TMRCAs for the antigenic gene segment were slightly smaller. Further, the difference in the mean TMRCAs for the antigenic and non-antigenic gene segment was smaller at higher mutations, most likely reflecting a concomitant increase in interference effects between the gene segments.

We also looked at the sensitivity of the coinfection rate by varying *β* from 0.0025 to 0.25 per day (Figure S7). When the frequency of the coinfected individuals was set at 1% in the total infected population (i.e *β* = 0.0025 per day) the TMRCA of the two gene segments was found to be very similar (Figure S7: mean difference in TMRCA is <0.5 years). While this level of coinfection corresponds well with empirical estimates (36-38), the difference in TMRCAs between antigenic and non-antigenic gene segments is not consistent with the observed evolutionary dynamics in Figure 2.

## DISCUSSION

We have developed a simple population genetic model to examine the role of non-antigenic gene segments in the adaptive evolution of seasonal influenza A viruses. In contrast to previous phylodynamic and predictive models of HA evolution, which have exclusively focused on HA (10, 12, 14, 15, 44), our approach allows us to evaluate the importance of selection on non-antigenic gene segments and intrasubtypic reassortment to the molecular and adaptive evolutionary dynamics of the virus genome. We find that the limited genetic diversity of non-antigenic gene segments and differences in TMRCAs between non-antigenic and antigenic gene segments are principally captured when selection on both antigenic and non-antigenic gene segments occurs and in the presence of low-level reassortment. Furthermore, while our results indicate that selection on the non-antigenic gene segment can slightly influence the evolutionary dynamics of the antigenic gene segment, reassortment increases viral adaptation in our model primarily by decreasing selective interference acting upon the non-antigenic gene segment, rather than the antigenic gene segment.

Given that only two segments are modeled in this study, it would be interesting to see if these results still hold when additional non-antigenic gene segments are considered. One prediction is that since linkage effects are expected to increase with additional gene segments, we are more likely to see the cumulative effect of selection acting on the non-antigenic segments on the antigenic gene segment. Furthermore, the fitness variation of each gene segment (and the overall virus) is also likely to increase, thus enabling selection to be more efficacious. Consequently, in light of this hypothesis, our finding that non-antigenic gene segment has minimal impact on the antigenic gene segment is likely to be overly conservative.

Since the coinfection level assumed in our model is ~5%, around 2.5% of the infected population is expected to carry a first-generation reassortant virus. Interestingly, this low level of reassortment is consistent with a recently estimated frequency of reassortment events observed among sampled virus genomes over time, at around 3.35% (25). Further evidence that the intrasubtypic reassortment is restricted at the between-host level comes from a recent finding that even at the within-host scale the effective reassortment rate is very limited (45). This indicates that the difference in the TMRCA across the seasonal influenza A virus genome is likely to arise from a low-level of reassortment in the virus population. Importantly, this has strong implications for the adaptive evolution of the virus, since it suggests that selective interference among gene segments has the potential to influence the fate of beneficial mutations in the genome.

Although reassortment is notoriously associated with pandemic influenza (46), there are several historical events in both seasonal influenza A/H3N2 and in seasonal influenza A/H1N1 where intrasubtypic reassortment has been implicated in antigenic cluster transitions (16, 22, 47). Furthermore, given that these instances are often associated with greater disease severity and incidence, akin to pandemic influenza, it also indicates that intrasubtypic reassortment can facilitate significant improvements in viral fitness. Consequently, this suggests that reassortment predominantly increases the rate of virus adaptive evolution by reducing selective interference effects across the genome.

We did not explicitly consider epistasis in our simulation model. There is evidence that epistatic interactions both within and between gene segments can drive the adaptive evolution of seasonal influenza A viruses. For example, T-cell immune escape mutations in NP have been enabled by stability-mediated epistasis (19, 20) and functional mismatches between the activities of the HA and the NA are known to decrease viral fitness considerably (48). However, to effectively model epistasis, a detailed knowledge about the fitness landscape of the virus genome, which is currently lacking, is necessary. Elucidating the epistatic interactions in influenza A viruses should be a focus of future work, since it could help explain the role that intrasubtypic reassortment plays in contributing to the adaptive evolution of seasonal influenza (49), and more broadly, it could help understand the epidemic (and even pandemic) potential of reassortant viruses.

Our findings that selection is likely to act upon both antigenic and non-antigenic gene segments and that reassortment can influence the rate of virus adaptive evolution have important implications for predicting future influenza strains. In particular, our study indicates that viral mutations are subjected to linkage effects within and to a somewhat lesser extent between gene segments, consistent with the conclusions of (37). As a consequence, we anticipate better forecasting can be achieved if the virus genetic background is considered as a whole, and is not just restricted to HA. This will be largely dependent on obtaining a more comprehensive understanding of the phenotypic variation in other gene segments, which we recommend should be a priority for future research.

